# In a Canine Model of Septic Shock, Cardiomyopathy Occurs Independent of Catecholamine Surges and Cardiac Microvascular Ischemia

**DOI:** 10.1101/2024.02.05.578927

**Authors:** Verity J. Ford, Willard N. Applefeld, Jeffrey Wang, Junfeng Sun, Steven B. Solomon, Harvey G. Klein, Jing Feng, Juan Lertora, Parizad Parizi-Torabi, Robert L. Danner, Michael A. Solomon, Marcus Y. Chen, Charles Natanson

## Abstract

**Background:** High levels of catecholamines are cardiotoxic and associated with stress-induced cardiomyopathies. Septic patients are routinely exposed to endogenously released and exogenously administered catecholamines, which may alter cardiac function and perfusion causing ischemia. Early during human septic shock, left ventricular ejection fraction (LVEF) decreases but normalizes in survivors over 7-10 days. Employing a septic shock model that reproduces these human septic cardiac findings, we investigated the effects of catecholamines on microcirculatory perfusion and cardiac function.

**Methods:** Purpose-bred beagles received intrabronchial *Staphylococcus aureus* (n=30) or saline (n=6) challenges and septic animals recieved either epinephrine (1mcg/kg/min, n=15) or saline (n=15) infusions from 4 to 44 hours. Serial cardiac magnetic resonance imaging (CMR), invasive hemodynamics and laboratory data including catecholamine levels and troponins were collected over 92 hours. Adenosine-stress perfusion CMR was performed on eight of the fifteen septic epinephrine, and eight of the fifteen septic saline animals. High-dose sedation was titrated for comfort and suppress endogenous catecholamine release.

**Results:** Catecholamine levels were largely within the normal range throughout the study in animals receiving an intrabronchial bacteria or saline challenge. However, septic *versus* non-septic animals developed significant worsening of LV; EF, strain, and -aortic coupling that was not explained by differences in afterload, preload, or heart rate. In septic animals that received epinephrine *versus* saline infusions, plasma epinephrine levels increased 800-fold, pulmonary and systemic pressures significantly increased, and cardiac edema decreased. Despite this, septic animals receiving epinephrine *versus* saline during and after infusions, had no significant further worsening of LV; EF, strain, or -aortic coupling. Animals receiving saline had a sepsis-induced increase in microcirculatory reserve without troponin elevations. In contrast, septic animals receiving epinephrine had blunted microcirculatory perfusion and elevated troponin levels that persisted for hours after the infusion stopped. During infusion, septic animals that received epinephrine *versus* saline had significantly greater lactate, creatinine, and alanine aminotransferase levels.

**Conclusions:** Cardiac dysfunction during sepsis is not primarily due to elevated endogenous or exogenous catecholamines nor is it principally due to decreased microvascular perfusion-induced ischemia. However, epinephrine itself has potentially harmful long lasting ischemic effects during sepsis including impaired microvascular perfusion that persists after stopping the infusion.

**Clinical Perspective:** *What is new?:*

- Myocardial depression of sepsis occurs without high levels of circulating catecholamines.
- Whereas large vessel coronary perfusion is known to be well maintained during sepsis, we show that during the myocardial depression of sepsis, in a model without exogenous catecholamine infusion, no perfusion abnormalities in the coronary microcirculation nor troponin elevations develop, indicating that the cardiac dysfunction of sepsis is not an ischemic injury.
- Epinephrine use during sepsis produces a form of injury tangential to the myocardial depression of sepsis.
- Epinephrine infusions depressed microcirculatory perfusion reserve and increased troponin I levels indicating a secondary prolonged mild ischemic effect on the myocardium.

*What are the clinical implications?:*

- Prolonged high doses of epinephrine can secondarily contribute to perfusion abnormalities.
- Decoupling the septic heart from microvascular perfusion abnormalities and ischemia may lead to better strategies for managing shock associated with severe infections. In clinical practice in septic patients particularly potentially with coronary artery disease, commonly used vasopressors that are less associated with increased lactate production than epinephrine, alongside adjunct cardiac microcirculatory vasodilators, could help better maintain or improve cardiac performance during septic shock.

## Introduction

During human septic shock, patients are exposed to high catecholamine levels from intrinsic release and exogenous administration. Independent of sepsis, high circulating catecholamine levels can have toxic effects on the heart contributing to an acute reversible heart failure syndrome commonly referred to as a stress-induced cardiomyopathy.^1, 2^ Sepsis, in both humans and animal models, also causes an acute reversible heart failure syndrome. In both septic humans and in animal models of sepsis, profound falls in left ventricular ejection fraction (LVEF) occur two days after the onset of shock (humans) or bacterial challenge (animal models) and reverse to near normal over 7-10 days.^3^ There is currently no consensus on the mechanism of this sepsis-induced cardiomyopathy; however, the impact of high levels of circulating endogenous and therapeutic exogenous catecholamines during septic shock remains a viable hypothesis.

In 2005, it was found in a study of critically ill patients with primarily non-cardiac diagnoses, 62% of patients who developed echocardiographic features of a stress-induced cardiomyopathy had sepsis.^4^ Further, a metanalysis of 23 separate case reports demonstrate an association between septic shock and stress-induced cardiomyopathy.^5^ More recently, observational studies have suggested that, in many cases, myocardial dysfunction in sepsis might be a stress-induced cardiomyopathy.^6, 7^ Sepsis can also result in a high catecholaminergic state in animal models due to intrinsic catecholamine release and extrinsic administration. In an awake murine cecal ligation model of sepsis where no exogenous catecholamines were administered, high levels of catecholamines were still detected due to endogenous release.^8^ In our previous large animal study of septic shock where both fluids and norepinephrine were titrated to physiologic end points, we found a strong negative correlation (−0.74, p = 0.04) in the first 24 hours after bacterial challenge between norepinephrine levels and LVEF.^9^ Catecholamine levels were found to be extremely high (on average 2000 pg/ml in animals in the first 24 hours after bacterial challenge) and the decreases in LVEF were profound (0.2 to 0.45 absolute percentage point drops).

Given the large body of supportive preclinical and clinical evidence suggesting that the myocardial depression of sepsis is a form of stress-induced cardiomyopathy, we decided to investigate this hypothesis.^5, 6^ Since, in patients with septic shock exhibiting life-threatening hypotension, it is neither possible nor ethical to withhold exogenous catecholamines or use sedatives and narcotics to suppress any stress-induced catecholamine response, we therefore utilized a canine model of sepsis which simulates the cardiovascular changes of human septic shock^5, 6^ to examine the impact of altering levels of catecholamines. In the published literature, epinephrine and isoproterenol have most commonly been used to create stress-induced cardiomyopathies in animal models, we chose epinephrine because it is also used clinically to treat septic shock.^10–13^ Finally, although human and animal sepsis studies have shown that large vessel coronary perfusion is not impaired during sepsis, the cardiac microcirculation has not been evaluated for impaired tissue perfusion as a cause of cardiac injury.^14, 15^ We previously found in our large animal model of sepsis-induced cardiac dysfunction that the coronary microcirculation is damaged, but this finding was confounded by the use of exogenous catecholamines.^10^ Electron microscopy (EM) demonstrated endothelial cell edema with a non-occlusive diffuse micro-vascular injury and fibrin deposition. A better insight into the role of catecholamines and microcirculatory tissue perfusion in sepsis-induced cardiomyopathy may generate novel approaches to managing the cardiomyopathy of septic shock.

## Methods

### Study Groups

Using a well-established canine model of bacterial pneumonia, we investigated the influence of exogenous and endogenous catecholamines and the role of microcirculatory perfusion on cardiac function during sepsis. Tracheostomized sedated, mechanically ventilated purpose-bred beagles (9 - 15 kg, 18 – 30 months, male, Marshall Farms) on day one at 0 hour (baseline) received either an intrabronchial challenge of *Staphylococcus aureus* (0.5 - 1.0 x10^9^ CFUs/kg) to induce sepsis or an equivalent intrabronchial inoculation volume of phosphate-buffered saline (PBS) as control.

### Animal Inclusion Criteria

The effects of catecholamines on myocardial function during sepsis were compared employing septic (n = 14) and non septic controls (n = 6). Seven of these animals received epinephrine and therefore were not used for the analysis in Figure 1 and 2. The other figures utilised the 14 septic animals that were paired each study week and received either an epinephrine (n=7) or saline infusion (n=7). We further conducted an experiment utilizing 16 septic animals that received an epinephrine (n=8) or saline infusion (n=8) who underwent a stress-adenosine CMR at baseline and 66h, and were analysed in Figure 4A-C and Figure 5A-C.

**Figure 1:**
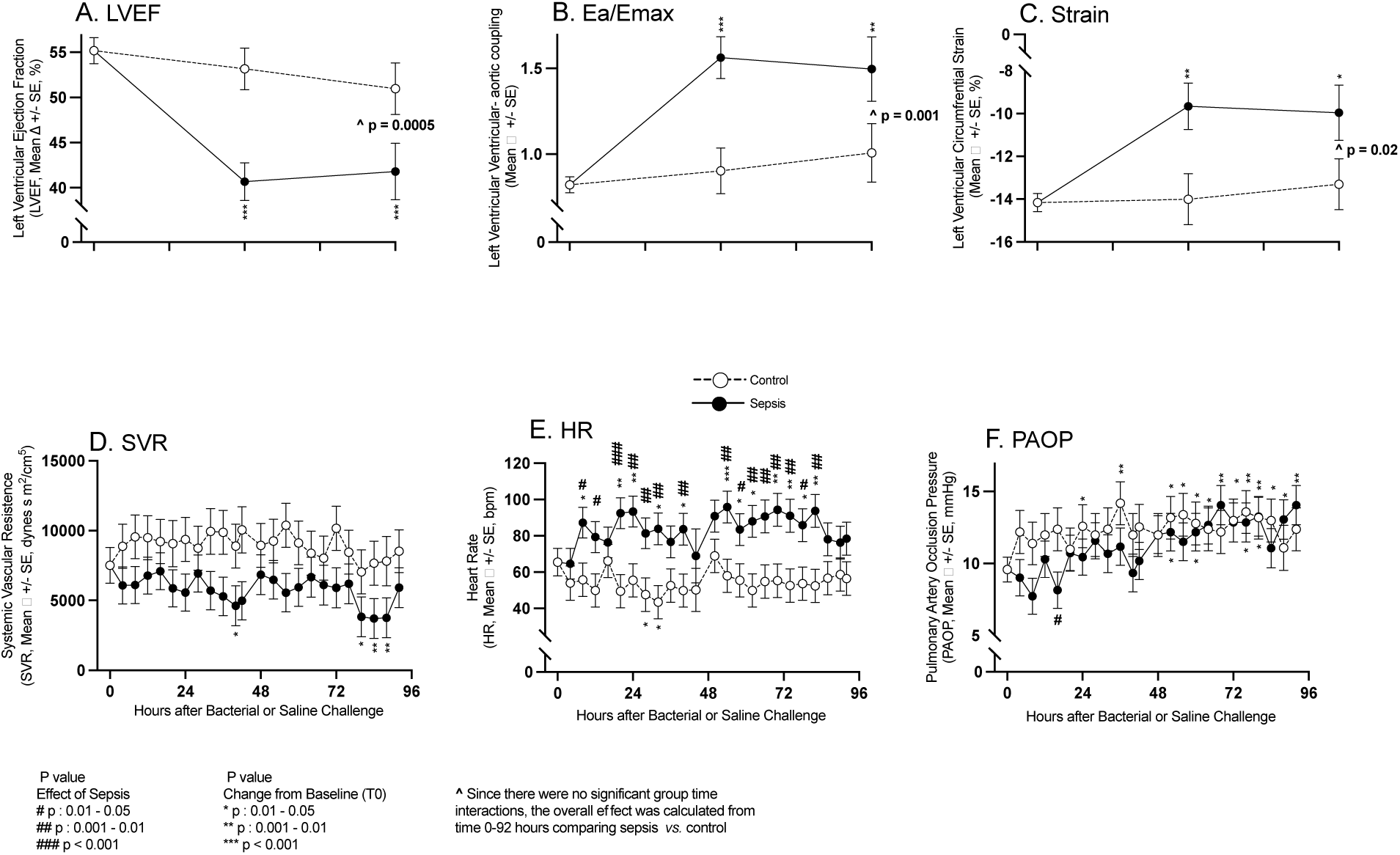
Serial CMR imaging and continuous hemodynamic monitoring parameters obtained in septic animals (closed circles) and non-septic controls (open circles). Serial mean changes from baseline to 96 hours are plotted from a common origin of the mean values of all animals at time 0 before intrabronchial bacterial or saline challenge. In the top panels are parameters (A-C) obtained by CMR imaging and in the bottom panels (D-F) are parameters obtained by invasive arterial and PACs.

**Figure 2:**
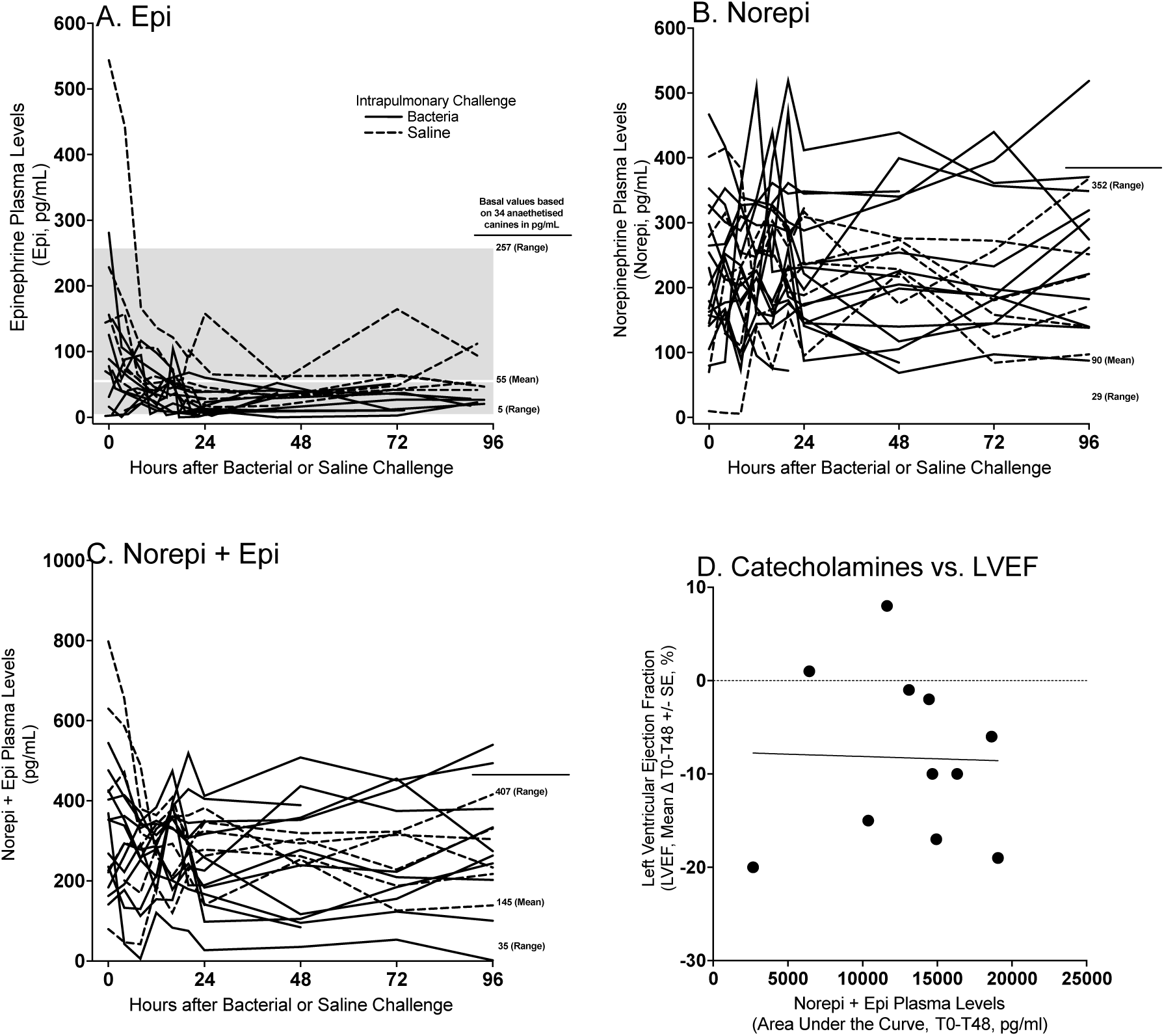
Serial individual animals plasma catecholamine levels as measured by ELISA in septic (filled line) or non-septic (dotted line) for epinephrine (Panel A), norepinephrine (Panel B) and norepinephrine and epinephrine combined (Panel C). Normal mean values and ranges for sedated canines for these catecholamines were obtained through the literature.^19^ In Panel D, individual animals’ changes from baseline to 48 hours in LVEF as measured by CMR were compared to the levels of endogenous catecholamines over 48 hours (AUC) for septic animals and controls.

Starting at 4 hours after bacterial challenge septic animals received a 40-hour continuous intravenous veterinary epinephrine infusion of 1 mcg/kg/min (Patterson Veterinary 1mg/mL) or saline equivalent. To maximize the potential for seeing the effects of catecholamine stress on the heart, we chose a supraphysiologic dose of epinephrine which is double the recommended intravenous infusion dose suggested for shock in canines.^16^ The epinephrine and saline infusions were terminated at 44 hours after intrabronchial bacterial or PBS challenge.

From the time of conclusion of experimental and control infusions (44 hours) until the end of the study (96 hours), the surviving animals continued to receive protocol-based treatment and monitoring with invasive hemodynamics and CMR to examine the effects of no exposure to exogenous catecholamines or prolonged exposure of high-dose catecholamines during sepsis on cardiac function and structure. At baseline and 62 hours after bacterial challenge, 16 septic animals, eight that received epinephrine and eight that did not receive epinephrine, underwent a CMR adenosine microcirculatory perfusion study. At 96 hours, all animals studied were deemed survivors and euthanized as per previously published protocols.^17^ All animals were treated equally, except for the experimental therapy and intrabronchial challenge.

### Animal Care

Animals were monitored and cared for by a clinician or trained technician around the clock for 96 hours to simulate patient care in a medical or animal hospital intensive care unit, as previously described.^17^ Throughout the study, animals received mechanical ventilation, sedation titrated to physiological endpoints, stress ulcer and venous thromboembolism prophylaxis, and their position was changed at set intervals to avoid stasis ulcers as previously described.^17^ All animals received daily intravenous ceftriaxone (50 mg/kg IV q24) starting 4 hours after bacterial or saline challenge until 96 hours or death. To avoid the influence of other exogenous catecholamines on cardiac function, no animal was administered vasoactive medications at any point during the study, except for those randomized to receive the continuous supraphysiologic epinephrine infusion. To examine cardiac function in as stress-free environment as possible, intravenous analgesia and sedation were targeted to eliminate any response to stimulation to minimize any endogenous catecholamine release. This was particularly important in the septic group that received saline to assess if cardiac depression during sepsis occurred even when the stress-induced catecholamine response was minimized. Maintenance fluids (2 ml/kg/h Normasol-M with 5% dextrose supplemented with KCl (27 mEq/l)) were administered to all animals starting at time 0 for 96h. A 20 ml/kg plasmaLyte bolus (Vetivex) was administered if pulmonary artery occlusion pressure (PAOP) fell below 10mmHg in a protocolized fashion until PAOP ≥ 10 mmHg was achieved. *Staphylococcus aureus* was prepared and administrated as in previous studies.^17^ The study protocol was reviewed and approved by the National Institutes of Health Clinical Center Institutional Animal Care and User Committee (CCM19-04 and CCM 22-04).

### Measurements

Before the above protocol was initiated, a tracheostomy was performed, an endotracheal tube placed, and femoral arterial and right heart thermodilution pulmonary artery catheters (PAC) were inserted under general anesthesia, as previously described.^17^ Femoral arterial catheters and PACs were used to perform serial invasive hemodynamic monitoring throughout the 96 hour duration of the study. These measurements included mean arterial pressure (MAP), central venous pressure (CVP), pulmonary artery systolic and diastolic pressures, mean pulmonary artery pressure (mPAP), PAOP, thermodilution derived cardiac output (CO), and heart rate (HR). Laboratory parameters were obtained from arterial blood gases, complete blood counts and serum chemistries (Heska, Loveland, CO). Endogenous plasma catecholamine levels (epinephrine, norepinephrine, and dopamine) were determined using commercially available canine ELISA kits (Life Diagnostics, West Chester, PA). Further, when the epinephrine infusion was running, the exogenous epinephrine values were determined using commercially available human ELISA kits (Abcam, Cambridge, UK). Troponin I was measured using a multidetector microplate reader (Synergy HT, BioTek Instruments, Winooski).

### Cardiac Magnetic Resonance Imaging

All animals underwent serial CMRs. All animals were transported to the scanner sedated, mechanically ventilated, and continuously monitored by a technician or clinician. Septic animals followed one of two time frames. Time frame one: A 3 Tesla MRI scanner (Philips Healthcare) acquired CMRs for 14 animals at baseline (T0), 42 hours after bacterial challenge (on epinephrine or saline infusion), and at the end of the study (96 hours, off infusion). Electrocardiogram-gated steady state free-precision cine and T2 images were acquired in mid-ventricular short axis and assessed for average plane values. Epicardial and endocardial contours were drawn on the short-axis slices at end-diastole and end-systole. A single perfusion rest scan with gadolinium gadobutrol 0.1 mmol/kg (Bayer Healthcare) followed by an adequate saline flush, and delayed enhancement scans were obtained. Time frame two: 16 animals, eight destined to receive an epinephrine infusion and eight a saline infusion, were imaged with CMR at baseline and at 62 hours after bacterial challenge (off infusion) as above and underwent a stress adenosine and rest microcirculatory perfusion study using the same gadolinium dose. Adenosine (140 mcg/kg/min) was administered as a continuous infusion for 5 minutes prior and then during the stress phase of the CMR perfusion study. The stress phase imaging preceded the rest phase imaging by approximately 10 minutes to allow washout of gadolinium contrast material and adenosine. All measurements for the CMRs were conducted using dedicated analysis software (NEOSOFT suiteHEART) by one of three investigators (VF, WA, JW) blinded to study animal treatment and each of the three-studies analysis checked for accuracy by a fourth investigator whose primary research is in CMR (MC), also blinded to study animal treatment group. Papillary muscles were included in the volumetric quantification of the LV.

### Statistical Analysis

Data was analyzed using linear mixed models to account for repeated measures and summarized as model estimate (standard error). We first tested the group-time interaction. If the interaction term was significant, groups were compared at each time point; otherwise, group comparisons were based on the main effects. Standard residual diagnostics were used to check model assumptions. All *p* values are two-sided and considered significant if *p* ≤ 0.05. For some variables, logarithm transformation was used when necessary. Statistical analysis (JS) was conducted using SAS version 9.4 (Cary, NC) with figure creation using GraphPad Prism 9.

## Results

### Myocardial Dysfunction During Sepsis

At 48 and 96 hours after bacterial challenge, septic animals had significant decreases in mean LVEF, and significant worsening of circumferential strain and ventricular-aortic coupling compared to both baseline and non-septic controls (Figure 1, Panel A-C), who had no significant changes in mean values in these same parameters throughout the study compared to baseline. In septic animals, at 40 and 80 to 88 hours, mean SVR was significantly decreased compared to baseline (Panel D). In non-septic controls, mean SVR over the entire 96-hour study was not significantly different compared to baseline and to septic animals. From 20 to 84 hours in septic animals, mean HR was significantly increased compared to baseline (Panel E). Septic animals had a significantly increased mean HR compared to their non-septic controls for the majority of timepoints (8 to 84 hours). At 52 to 92 hours in septic animals, there was a significant increase in mean PAOP compared to baseline (Panel F). At 24 to 84 hours after PBS challenge in non-septic animals, mean PAOP was significantly increased compared to baseline. At only one time point (16 hours) there was a significant decrease in the mean PAOP in septic animals compared to non-septic controls. From 0 to 96 hours, septic animals *versus* non-septic controls had no significant differences in the quantity of total fluids received (158 ml/kg +/- 18, *vs.,* 156 ml/kg +/- 19 respectively). There were no significant differences in afterload or preload between control and septic animals to explain the cardiac depression seen in septic animals. The increases in HR should increase inotropy and decrease cardiac filling, therefore increasing LVEF.^18^ However, the opposite was found to be true. Thus, hemodynamic changes cannot account for the profound myocardial depression. Next, we examined whether the decline in LVEF found here was associated with catecholamine level elevations.

### Plasma Catecholamines Levels in Sedated Septic Animals

At T0, when animals were transferred from the surgical suite after undergoing general anesthesia to intensive care to start the study sedation protocol, several animals had baseline elevations in catecholamine levels before the sedation protocol was fully initiated (i.e., perfectly dosed). After bacterial challenge (T0), in the absence of exogenous catecholamine administration, serial plasma epinephrine, norepinephrine, and combined catecholamine levels were within or slightly above the normal range for sedated otherwise healthy canines (Figure 2, Panel A-C).^19^ Specifically, the only elevations in catecholamine levels above the normal range occurred in the plasma norepinephrine concentration of septic animals. In these animals, the elevations occurred transiently and were only minimally above the upper limit of normal (Figure 2, Panel B) for sedated animals. There were no marked differences between septic animals and controls in serial catecholamine levels from baseline to 96 hours. The serial dopamine levels were also measured between septic animals and non-septic controls but normal ranges for sedated canines were not available. However the plasma dopamine levels over 96h in septic *versus* non septic controls were not higher (data not shown). There was no significant association in septic animals between changes in LVEF from 0 to 48 hours, the time of maximum decrease in LVEF, and the area under the curve for combined catecholamine (norepinephrine + epinephrine) levels from 0 to 48 hours (Panel D, p=0.94). Therefore, no associations were seen between LVEF depressions and endogenous catecholamine levels.

### The Cardiovascular Effects of IV Epinephrine During Infusion (4-44 hours)

During the 40-hour control infusion of saline in septic animals, significant decreases in mean MAP occurred from 24 to 42 hours after bacterial challenge compared to baseline (Figure 3, Panel A). In contrast, in septic animals who received a 40-hour infusion of epinephrine, significant marked increases were seen in mean MAP from baseline during the infusion at 16 to 42 hours. Septic animals receiving the epinephrine infusion had significant increases in MAP compared to septic animals who received saline infusions during most of the later infusion times (16-to-42-hour timepoints). From 0 to 42 hours, septic animals that received epinephrine had an overall marked significant increase in mean SVR during the 40-hour infusion compared with those septic animals who received a saline infusion (Figure 3, Panel B). Consistent with the development of clinical pneumonia, all septic animals, both the epinephrine and saline infusion groups, developed significant increases in mPAP from baseline at 16h to 42h (Panel C). From 8 to 36 hours, septic animals who received epinephrine had a significantly greater increase in mPAP from baseline compared to septic animals who received a saline infusion. In septic animals receiving an epinephrine infusion and in those receiving the saline infusion, mean HR was significantly elevated from baseline at multiple time points between 8 to 42 hours.

**Figure 3:**
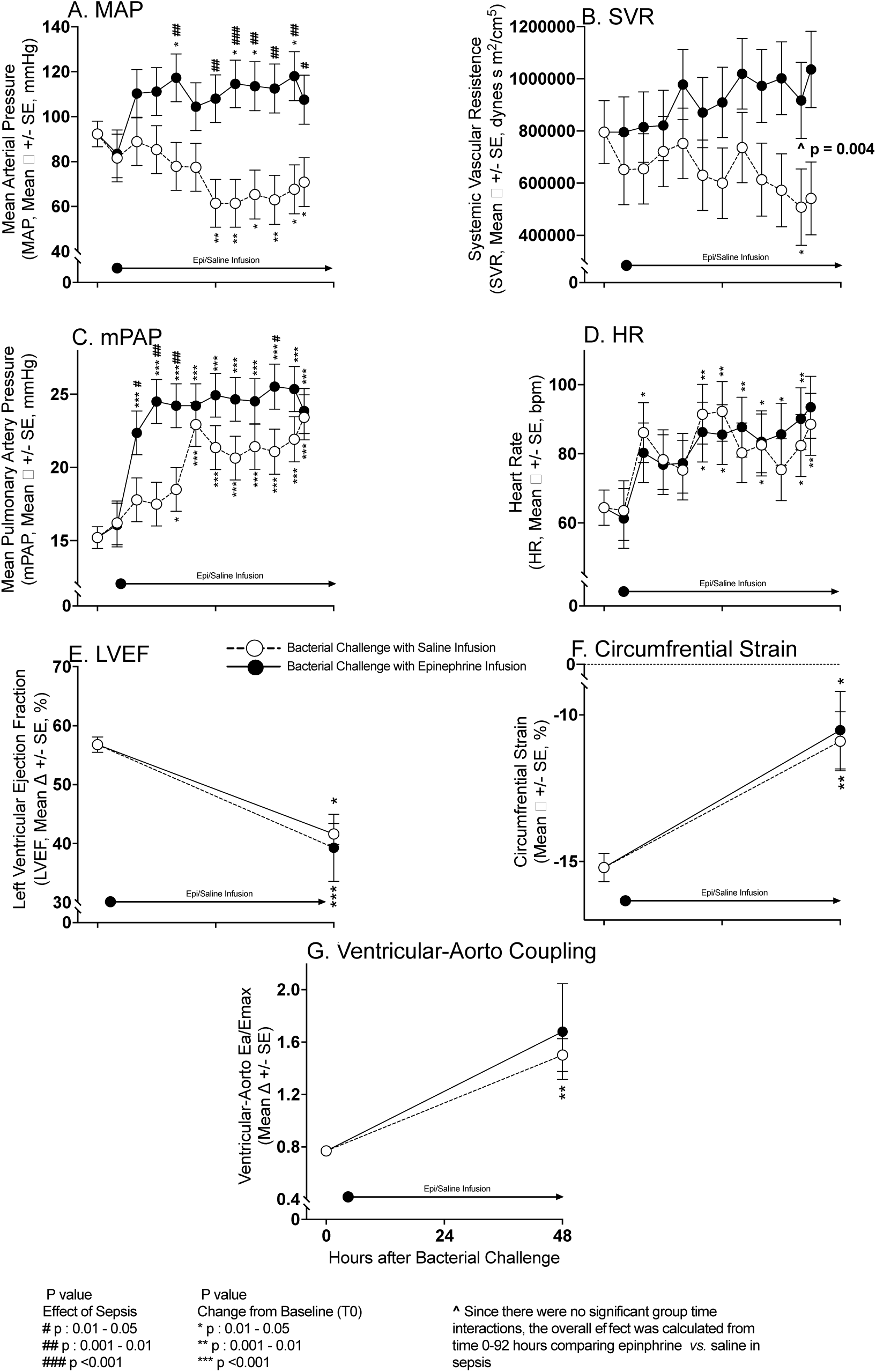
Serial continuous hemodynamic monitoring and CMR Imaging during experimental septic shock. Serial mean hemodynamic changes ascertained by arterial and PAC measures from baseline to 44 hours in septic animals while on epinephrine (filled circles) or saline Infusions (open circles) in Panels A-D. These changes are plotted from a common origin of the mean values of all septic animals at time 0. Serial mean cardiac function measures as obtained by CMR imaging from time 0 to 44 hours on epinephrine and saline infusions in Panels E-G.

However, there were no significant differences in mean HR from baseline between septic animals receiving epinephrine and those receiving saline at any time during the infusion (Panel D). Septic animals receiving epinephrine *versus* saline had similar significant decreases in mean LVEF at 48 hours compared to baseline and similar significant worsening of mean circumferential strain compared to baseline on CMR (Panel E-F). At 48 hours, compared to baseline, no significant difference was found in the worsening of ventricular-aortic coupling between septic animals receiving epinephrine and those receiving saline infusions (Panel G). In septic animals that received a 40-hour epinephrine infusion, at 24 hours (mid infusion), mean epinephrine levels were 17,928.48 ± 8,334 pg/ml and in those receiving saline at 24 hours mid infusion mean epinephrine levels were 22.1 ± 6.5 pg/ml. Therefore, epinephrine infusions during sepsis increased epinephrine levels 800-fold and greatly increased afterload in the pulmonary and systemic circuits (as evidenced by marked significant increases in mPAP, SVR, and MAP). However, septic animals receiving epinephrine *versus* saline, during and after infusions, had no significant further worsening of LV; EF, strain, or -aortic coupling.

### The Cardiovascular Effects of IV Epinephrine 2 Days After Discontinuation of the Infusion (46-92 hours)

Septic animals who received a 40-hour saline infusion (between 4 to 44 hours after bacterial challenge) had a significant decrease in mean LVEF compared to baseline at the 66- and 92-hour time points (approximately 22 and 48 hours after conclusion of the saline infusion) (Figure 4, Panel A). Comparing septic animals receiving epinephrine or saline infusions, there were no significant differences in these mean LVEF decreases from baseline throughout the study. Worsening of circumferential strain compared to baseline at 66 and 92 hours (Figure 4 Panel B) was similar in septic animals that received epinephrine and those that received saline infusions. Lastly, changes from baseline in septic animals that received epinephrine or saline infusions did not differ for ventricular-aortic coupling throughout. Therefore, no significant late differences at 66 and 96 hours were found in markers of cardiac function (LV; EF, strain, -aortic coupling) between septic animals that had received saline or those that had received epinephrine. These time points, 66 and 92 hours occurred 22 and 48 hours respectively after conclusion of the infusions (Figure 4 Panel A, B and C). At timepoints when the CMRs were performed (66 and 92 hours), no significant differences were seen in mean SVR, mean HR, and PAOP (Figure 4 Panels D, E and F, respectively) between septic animals that had received either epinephrine or saline infusions. After discontinuation of infusions at 44 hours, from 45-96 hours after bacterial challenge, there were also no marked differences between septic animals who had received epinephrine and those who had received saline in serial plasma levels of plasma epinephrine (Panel G), norepinephrine (Panel H), and dopamine (Panel I). Thus, while the epinephrine infusion was running, there were dramatic effects on hemodynamics and catecholamine levels but no worsening of myocardial dysfunction. Moreover, after stopping the infusion, there were no residual significant hemodynamics effects or worsening of myocardial depression for two days afterwards.

**Figure 4:**
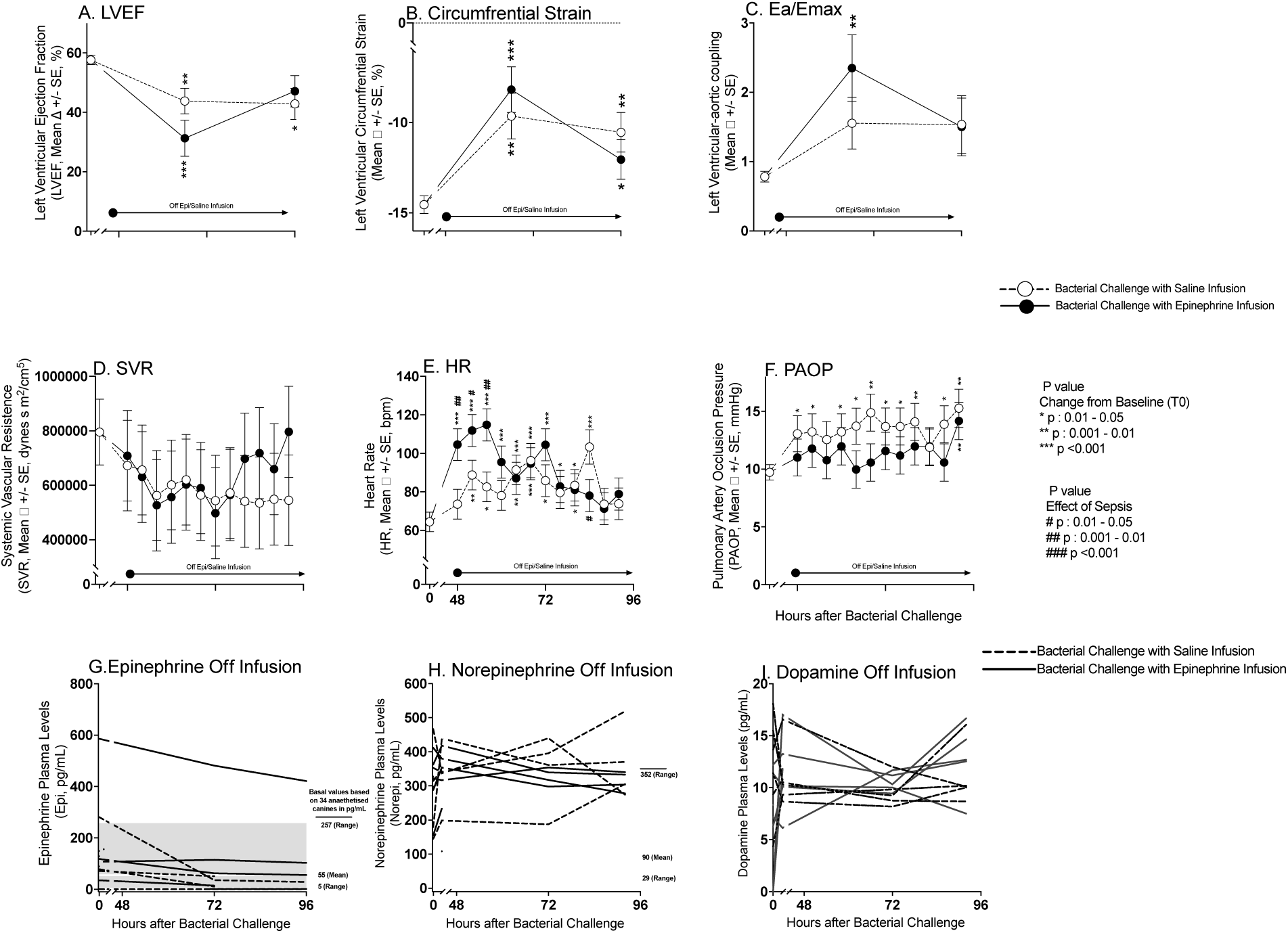
Serial CMR imaging and continuous hemodynamic monitoring during experimental septic shock after discontinuation of epinephrine infusions. Cardiac function changes obtained from CMR from baseline to 62 and 96 hours are plotted by closed circles for septic animals that received epinephrine infusions and open circles for septic animals that received saline infusions from a common origin of the mean values of all animals before intrabronchial challenge. Continuous hemodynamic measure changes obtained by arterial, and PACs are shown from baseline to 48 and 96 hours in Panel D to F. Serial individual animals plasma catecholamines levels, as measured using ELISA, from 48 to 96 hours (Panel G-I). Epinephrine, norepinephrine and dopamine levels shown by dotted lines for all septic animals that received saline prior, and continuous lines for animals that received epinephrine.

### Effect During Sepsis of Prior Epinephrine Infusion on Myocardial Microcirculatory Perfusion Reserve and Troponin Levels

At baseline, prior to the onset of sepsis and initiation of epinephrine/saline infusions, the uptake of myocardial gadolinium during the adenosine stress perfusion CMR was similar in animals destined to receive epinephrine and those bound to receive saline infusions (Figure 5, Panel A). At baseline, five minutes after discontinuing adenosine (rest perfusion), the uptake of gadolinium in the myocardium decreased similarly in all these animals. The difference between gadolinium uptake during adenosine infusion and the rest measurement represents normal microcirculatory perfusion reserve at baseline. At 62 hours, myocardial gadolinium uptake markedly increased during the adenosine infusion in septic animals that had received saline infusions 22 hours before (first open bar, Panel B) compared to gadolinium uptake at baseline in those same animals (first open bar, Panel A). In contrast, at the same late timepoint (62 hours), septic animals that had received an epinephrine infusion 22 hours before had decreased myocardial gadolinium uptake (first filled bar, Panel B) compared to baseline (first filled bar, Panel A). During the rest perfusion scan performed at 62-hour time point, both septic animals that received either epinephrine or saline infusions had no difference in myocardial perfusion compared to baseline. Consequently, the effect of sepsis on microcirculatory perfusion reserve is significantly different and opposite depending on whether an animal had received a prior infusion of epinephrine, even though the epinephrine infusion had been discontinued almost one day ago. Microcirculatory perfusion reserve (stress – rest) of the myocardium was significantly increased by sepsis in the absence of exogenous epinephrine (Panel C). Epinephrine blunted this vasoreactivity at 62 hours, decreasing the microcirculatory perfusion reserve of cardiac tissue in the septic animals 22 hours after discontinuation of the epinephrine infusion. This suggests an effect that is not mediated by direct action of epinephrine at its receptor.

**Figure 5:**
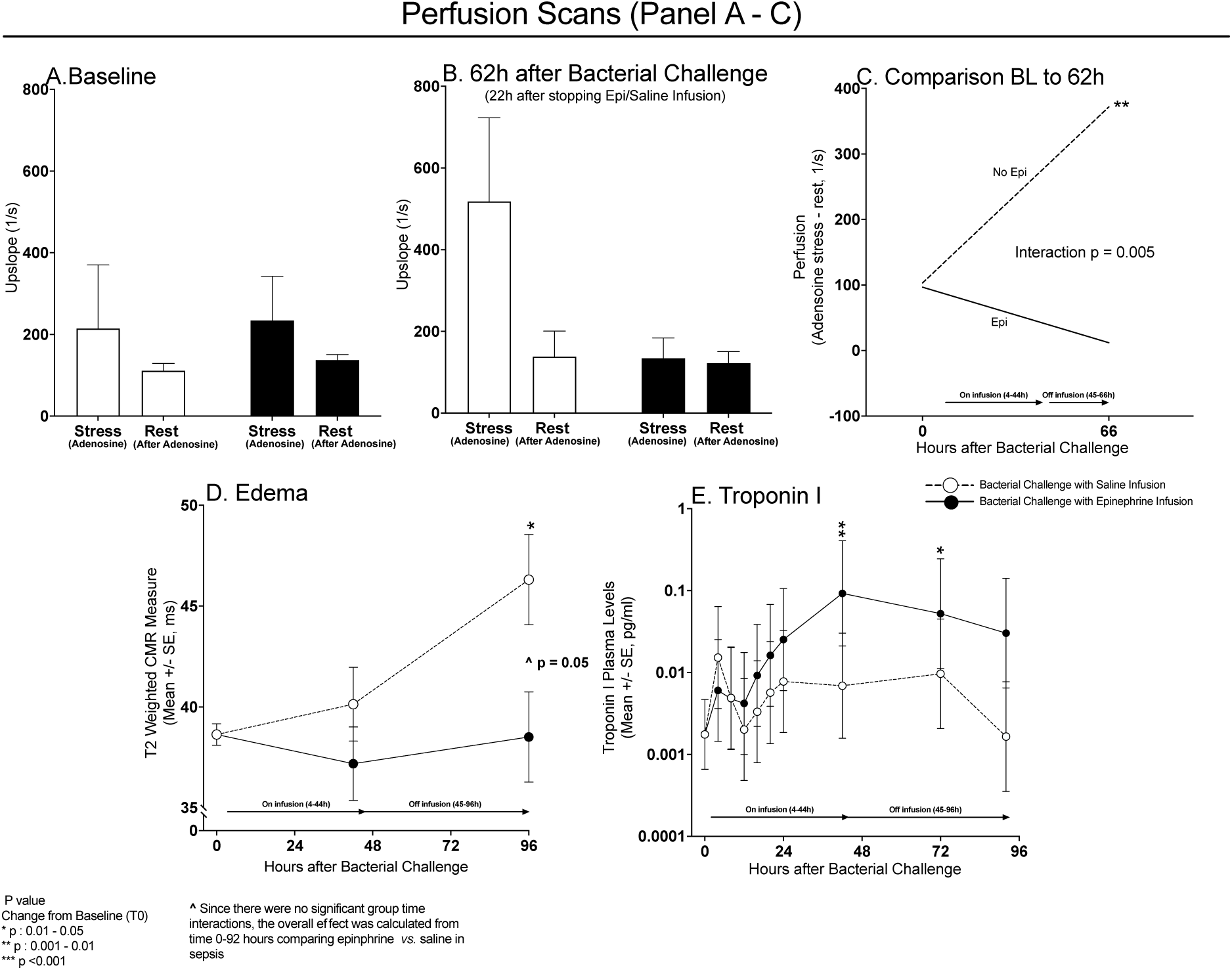
CMR derived adenosine-stress and rest perfusion scans at baseline (Panel A) and 66h (Panel B) after bacterial challenge for septic animals receiving a 40 hours continuous epinephrine infusion (filled bars) or saline infusions (open bars) starting at 4 hours after bacterial challenge. The change from baseline to 62 hours in perfusion (adenosine stress – rest) is shown in Panel C for septic animals that did not receive epinephrine (dotted line) and septic animals that received epinephrine (continuous line). There is a quantitative interaction with prior epinephrine *vs.* saline infusions significantly reducing microcirculatory perfusion reserve in septic animals at 62 hours. Mean CMR derived T2 measures (edema, Panel D) before bacterial challenge (0 hours), at 44 hours (on epinephrine or saline infusion), and 52 hours after epinephrine or saline infusions ended (or 96 hours after bacterial challenge) in septic animals receiving epinephrine (filled circles) or saline (open circles). Panel E, serial mean changes in plasma troponin I levels plotted from a common origin (time 0 hours values in all animals) comparing septic animals receiving epinephrine (closed circles) or saline infusion (open circles).

**Figure 6:**
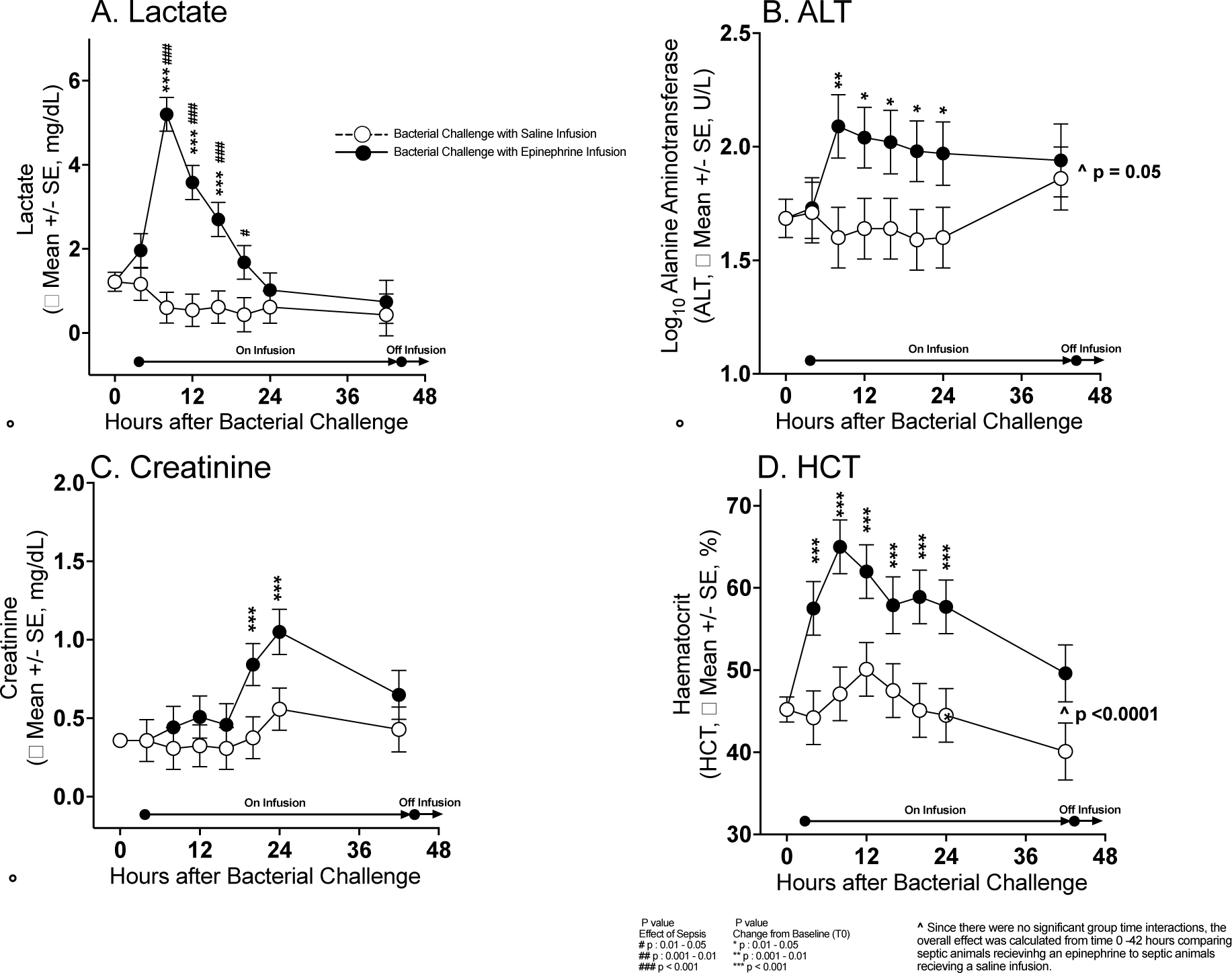
Serial mean change from baseline (before bacterial challenge at 0 hours) to 48 hours in lactate, ALT, creatinine and HCT levels plotted from a common origin (all animals’ mean values at time 0 hours) comparing septic animals receiving a 40 hours epinephrine (filled circles) or saline infusion (open circles).

Therefore, sepsis markedly increases microcirculatory perfusion reserve and epinephrine during sepsis induces a loss of myocardial microcirculatory perfusion reserve that persists long after discontinuation of the drug (Qualitative interaction, p = 0.005).

In septic animals not receiving epinephrine, there was a 20% increase (+ 7.68 ± 2.24 ms, p = 0.01) in LV wall edema from baseline to 96 hours (Panel D). The epinephrine infusions significantly decreased this edema. Throughout the 96-hour study, there were no significant elevations in mean troponin I level from baseline in control animals that had not received an epinephrine infusion (Panel E). In septic animals receiving an infusion of epinephrine, troponin I levels were significantly elevated from baseline at 42 and 72 hours, consistent with the epinephrine-induced tissue perfusion abnormalities. As stress CMR imaging at baseline confirmed absence of flow limiting epicardial coronary disease, the perfusion abnormality at late time points in septic canines exposed to a prolonged epinephrine infusion indicates the development of microvascular dysfunction. A prolonged infusion of epinephrine during sepsis interfered with the increase in microcirculatory perfusion reserve and was associated with a troponin I leak.

### Effects of Epinephrine Infusions on Other Organs

Septic animals receiving epinephrine infusions from 4 to 44 hours had significantly raised mean lactate levels compared to baseline at 8 to 24 hours and compared to septic controls at 8 to 20 hours. Mean alanine aminotransferase and hematocrit levels were increased throughout the epinephrine infusion compared to saline controls. Creatinine levels were also elevated compared to baseline from 20 to 44 hours in animals receiving epinephrine. After stopping epinephrine, no significant elevations in any of the above parameters was seen (data not shown). Thus, there is marked multiorgan vasoconstriction during infusions of high-dose epinephrine causing transient ischemia and short-term abnormalities in liver and renal function that reverse immediately after the infusion is stopped.

### Other Laboratory Values

There were isolated significant findings comparing mean serial values for septic animals who received an epinephrine infusion compared to septic animals who received a saline infusion for serum cytokines, chemistries, complete blood count, electrolytes, and arterial blood gas parameters; none of these isolated differences explain our cardiac findings. These results are available in an e-supplementary Results.

## Discussion

In this model of the myocardial depression of sepsis, no exogenous catecholamines were administered in septic control animals receiving a saline infusion and non-septic animals that did not receive a bacterial challenge. Further in these septic and non-septic controls endogenous catecholamine release was blunted with analgesia and sedation. In these septic and non-septic controls catecholamine levels remained within, or very near, the normal range for anesthetized canines throughout the study.^19^ However, we found that myocardial depression still occurred. In septic animals compared to non septic controls, profound highly significant drops in LVEF and significant worsening of circumferential strain and ventricular-aortic coupling occurred over two days, which occurred in the absence of increased endogenous and exogenous catecholamines. These findings are not explained by changes in afterload, preload, or heart rate. Therefore, in this study normal or low levels of catecholamines do not prevent the pattern of cardiac dysfunction commonly seen in sepsis. This suggests that the myocardial dysfunction of sepsis is not primarily a catecholamine mediated process where high levels produce a stress-induced cardiomyopathy.

Certain inotropes are known to cause significant negative effects on survival in the heart failure literature.^20–22^ To establish the impact of high levels of catecholamines on cardiac function during sepsis we examined if the decreases in cardiac function associated with sepsis was worsened by pharmacologically elevated catecholamine levels. Sedated septic animals with cardiac dysfunction received a supraphysiologic high-dose epinephrine infusion for 40 hours. This infusion increased plasma epinephrine levels 800-fold (up to 20,000 pg/ml). These high doses of epinephrine caused substantial increases in systemic pressures in septic animals; however, there was no observed worsening of sepsis-induced changes in cardiac function including LVEF, ventricular-aortic coupling, and left ventricular circumferential strain measurements during infusion or for approximately two days afterwards. Therefore, challenges with supraphysiologic doses of catecholamines as well as suppressing endogenous catecholamine release to near normal levels does not measurably worsen the cardiac depression of sepsis. This further affirms that the cardiac depression of sepsis is not primarily a stress-induced cardiomyopathy and represents some separate pathophysiologic entity.

Epinephrine was associated with cardiac microvascular non-occlusive abnormalities, however, the myocardial injury attributable to epinephrine infusion was distinct from the myocardial depression observed in sepsis. For two days after discontinuation of epinephrine infusions, there was evidence of mild ischemia with significant reduction of microcirculatory reserve and minimal but significant troponin I level elevations compared to baseline. These findings were not seen in septic animals that did not receive epinephrine and represent a sustained late-occurring specific effect on microcirculatory perfusion particular to high-dose epinephrine. In addition to increased lactate levels associated with epinephrine infusions, liver enzymes were elevated, and renal function was reduced. In these other organs, unlike the heart, these abnormalities completely reversed after termination of the infusion. Further, in contrast to other organs and despite decreased microvascular perfusion, during the epinephrine infusion the heart showed no overt signs of injury, i.e., cardiac edema was decreased, and no troponin leak was detected. This observation suggests that by reducing coronary capillary perfusion, epinephrine did have directly measurable acute effects on the coronary microcirculation. Once epinephrine is stopped, perfusion returned toward normal, and troponins were then released into the circulation.

In this animal model of the cardiac depression of human sepsis we found independent of catecholamines, there is a resultant clinically important suppression of LVEF. The accompanying increase in microcirculatory perfusion reserve during sepsis indicates that myocardial depression in sepsis is not caused by inadequate tissue perfusion. These findings combined with the lack of troponin I elevation in the absence of epinephrine infusions exclude microcirculatory ischemia and decreased tissue perfusion as necessary conditions for the development of the cardiac depression of sepsis.

In the 1980s, the experimental use of coronary sinus venous catheterization in human sepsis demonstrated that the myocardial dysfunction during sepsis was not associated with reductions in coronary blood flow^15, 23^ or increased lactate levels despite profound decreases in LVEF. Further suggesting that microcirculatory ischemia was not present. The absence of a sepsis-induced decrease in coronary perfusion as a cause of myocardial dysfunction was further bolstered by our peritonitis large animal model, where direct coronary artery flow probes were used to demonstrate normal or increased coronary flow despite development of profound myocardial depression.^14^ This current study adds to prior findings by demonstrating that sepsis increases microcirculatory reserve and sepsis alone does not necessarily elevate troponin levels despite profound LVEF depression. These data effectively rule out microvascular ischemia and inadequate tissue perfusion as a primary cause of sepsis-induced myocardial depression.

It is unclear why sepsis results in, not just a preservation, but an increase in microcirculatory reserve. During the myocardial depression of sepsis, the non-occlusive edematous microcirculatory injury occurring predominantly in endothelial cells, but also the interstitium and myocytes, may cause a compensatory increase in nitric oxide (NO) sensitivity or release at the microvascular level.^24^ The administration of high-dose vasopressor infusion may substantially alter the microcirculation in sepsis resulting in further endothelial and downstream tissue injury preventing NO release, or causing depletion of NO reserves which decreases microcirculatory perfusion reserve. The decreased microcirculatory reserve with elevated troponin I suggests that high-dose catecholamines produces downstream myocardial tissue ischemia due to imbalances in the supply of/demand for coronary blood flow. Whereas myocardial tissue ischemia may well contribute to cardiac dysfunction in some patients, our data suggests that it is not the primary cause of sepsis-induced myocardial depression.

### Limitations

There are limitations to the interpretability of our findings. Different doses of sedatives, narcotics, and epinephrine infusions may have altered our findings. Similarly, lower dose epinephrine or different vasopressor agents may result in different findings. Nevertheless, the use of sedation to suppress endogenous catecholamine release and supraphysiologic epinephrine infusions demonstrate that high levels of catecholamines are not necessary to induce the myocardial depression of sepsis. We studied young canines with no underlying diseases and in clinical practice, patients with sepsis have a myriad of comorbidities including epicardial coronary disease and underlying microvascular dysfunction. Therefore, human subjects may demonstrate different physiologic responses. However, the changes in cardiac function are remarkably similar in our model to what is observed in humans, suggesting this may be the highly conserved mammalian response to severe systemic infection.

## Conclusions

We demonstrate here that sepsis-induced myocardial depression is not primarily a catecholamine-induced cardiomyopathy and does not arise from cardiac microcirculatory abnormalities that cause tissue ischemia. These studies add to our understanding of the pathophysiology of cardiac dysfunction in septic shock. Further, sepsis-induced myocardial depression is associated with myocardial edema which is unrelated to catecholamine toxicity and/or tissue ischemia. In survivors, this is rapidly reversed over 7 to 10 days by the removal and repair of presumably damaged cellular components (manuscript submitted, Circulation). The precise mechanistic relationship of edema formation to global cardiac dysfunction during sepsis remains to be elucidated. It is possible in some clinical situations that elevated catecholamines levels and microvascular tissue ischemia contribute to the increased severity of the cardiac dysfunction of sepsis. However, we show here that this is not the primary etiology. Edema, tissue injury, and subsequent repair mechanisms remain fertile targets for future sepsis and cardiovascular research.

## Supporting information

Supplemental results and figures

## SOURCES OF FUNDING

This work was supported by NIH intramural funding from the NIH Clinical Center.

## Role of funding source

The work by the authors was conducted as part of US government– funded research; however, the opinions expressed are not necessarily those of the National Institutes of Health (NIH).

## DISCLOSURES

The authors do not have any conflicts to disclose

## SUPPLEMENTAL MATERIAL

Results

Figure S1-S5

References 19

## Non-standard Abbreviations and Acronyms

ABGs: Arterial Blood Gas
BNP: Brain Naturietic Peptide
CBCs: Complete Blood Count
CMR: Cardiac Magnetic Resonance
CVP: Central Venous Pressure
EDV: End Diastolic Volume
EM: Electronic Microscopy
HR: Heart Rate
LV: Left Ventricular
LVEDV: Left Ventricular End Diastolic Volume
LVEF: Left Ventricular Ejection Fraction
MAP: Mean Arterial Pressure
PBS: Phosphate Buffer Solution
PACs: Pulmonary Artery catheters
PAOP: Pulmonary Artery Occlusion Pressure
PAP: Pulmonary Artery Pressure
PVR: Pulmonary Vascular Resistence
SV: Stroke Volume
SVR: Systemic Vascular Resistance

## Notes

### Competing Interest Statement

The authors have declared no competing interest.

