## Supplemental results and figures for "In a Canine Model of Septic Shock, Cardiomyopathy Occurs Independent of Catecholamine Surges and Cardiac Microvascular Ischemia"

**Supplementary Laboratory Results**

**Cytokines (IL-6, -8 -12 and -10, interferon- γ, von Willebrand factor, Tumor necrosis factor- α, p-selectin, and Monocyte Chemoattractant Protein)**

There were no significant differences in any cytokine measure comparing septic animals that received an epinephrine *vs.* saline infusion (e-supplementary figure 1). There were significant increases in septic animals who received epinephrine or saline infusion, compared to baseline, in mean IL-6, IL-8 and MCP. There were no significant differences compared to baseline in septic animals who received epinephrine or saline infusion in mean IL-12, TNF-α , and p-selectin levels. Overall, there were no significant differences compared to baseline in mean IL-10, IFN-γ, and vWF levels in both septic animals that receive epinephrine or saline infusion.

**Serum Chemistries, Complete Blood Count, Arterial Blood Gases, and Electrolytes**

Mean pH in septic animals receiving epinephrine was significantly less at 8 and 12 hours (e-supplementary figure 2, Panel A) compared to septic animals receiving saline. Mean $pCO_2$ was significantly less in septic animals receiving epinephrine compared to saline controls at 24 and 44 hours (Panel B). There were significant changes in septic animals who received epinephrine or saline infusion, compared to baseline, in mean arterial pH, $pCO_2$ and $pO_2$ levels throughout.

There were no significant differences in change from baseline for mean total protein, BUN, and albumin levels in septic animals receiving epinephrine *vs.* saline (e-supplementary figure 3). There were isolated significant, but not clinically relevant, increases in mean serum Total Protein and BUN in septic animals that received epinephrine or saline (Panel A and B).

There were significant decreases compared to baseline in mean serum albumin levels (Panel C) and mean BUN/Cre (Panel D) in both septic animals receiving epinephrine or saline. Mean BUN/Cre was significantly less in septic animals receiving epinephrine compared to saline.

Septic animals that received epinephrine had significantly increased mean glucose and decreased mean serum potassium levels compared to septic animals that received saline (e-supplementary figure 4). Further, septic animals that received epinephrine had significant differences compared to baseline in mean serum glucose (Panel A) and potassium (Panel B) whilst septic animals receiving saline only had differences in mean potassium only. There were no significant changes from baseline or significant differences in septic animals that received epinephrine or saline for mean serum sodium or chloride levels (Panel C and D, respectively).

There were no other significant differences between septic animals receiving epinephrine or saline infusion in mean lymphocyte counts throughout the study (e-supplementary figure 5, Panel A). However, there were significant overall differences between septic animals receiving epinephrine compared to saline in mean hemoglobin, platelet, and eosinophil counts whilst on infusion (Panel B-D). Septic animals receiving epinephrine had significantly higher mean WBC counts at 20 to 44 hours compared to septic animals receiving saline (Panel E). There were significant changes from baseline in both septic animals receiving epinephrine or saline in mean lymphocytes, hemoglobin, platelets, eosinophils and WBC counts throughout the study.

### Figures

#### e-supplementary figure 1

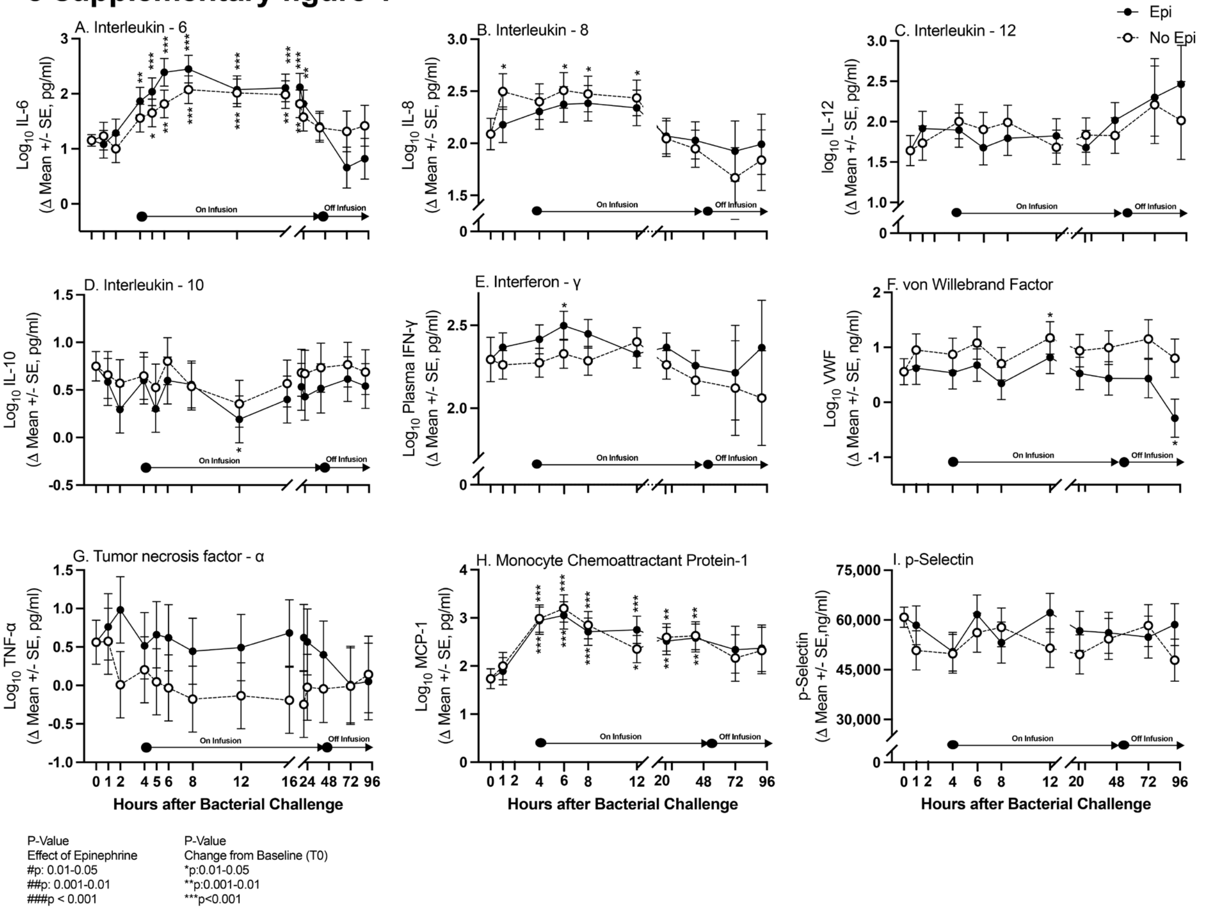


**e-supplementary figure 1: The format is similar to figure 1**

#### e-supplementary figure 2

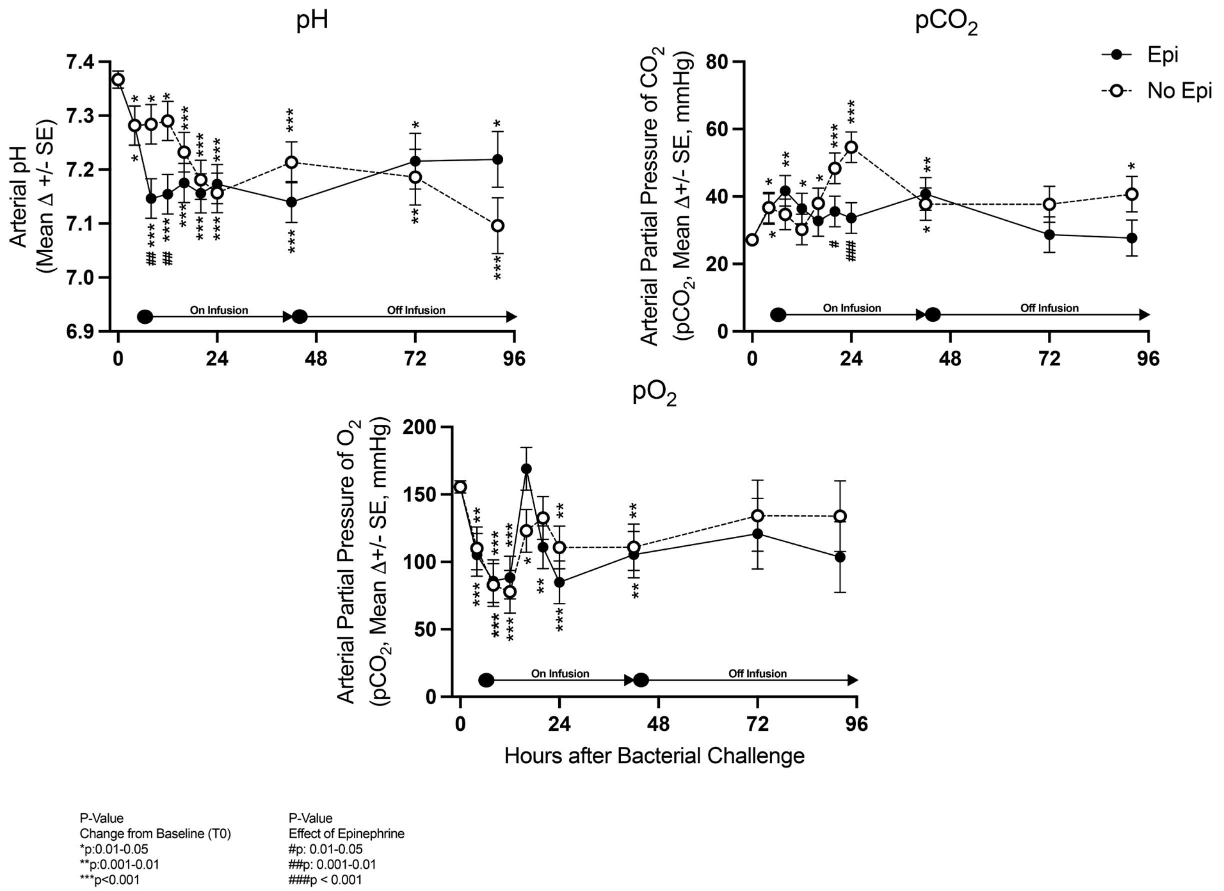


**e-supplementary figure 2: The format is similar to figure 1**

#### e-supplementary figure 3

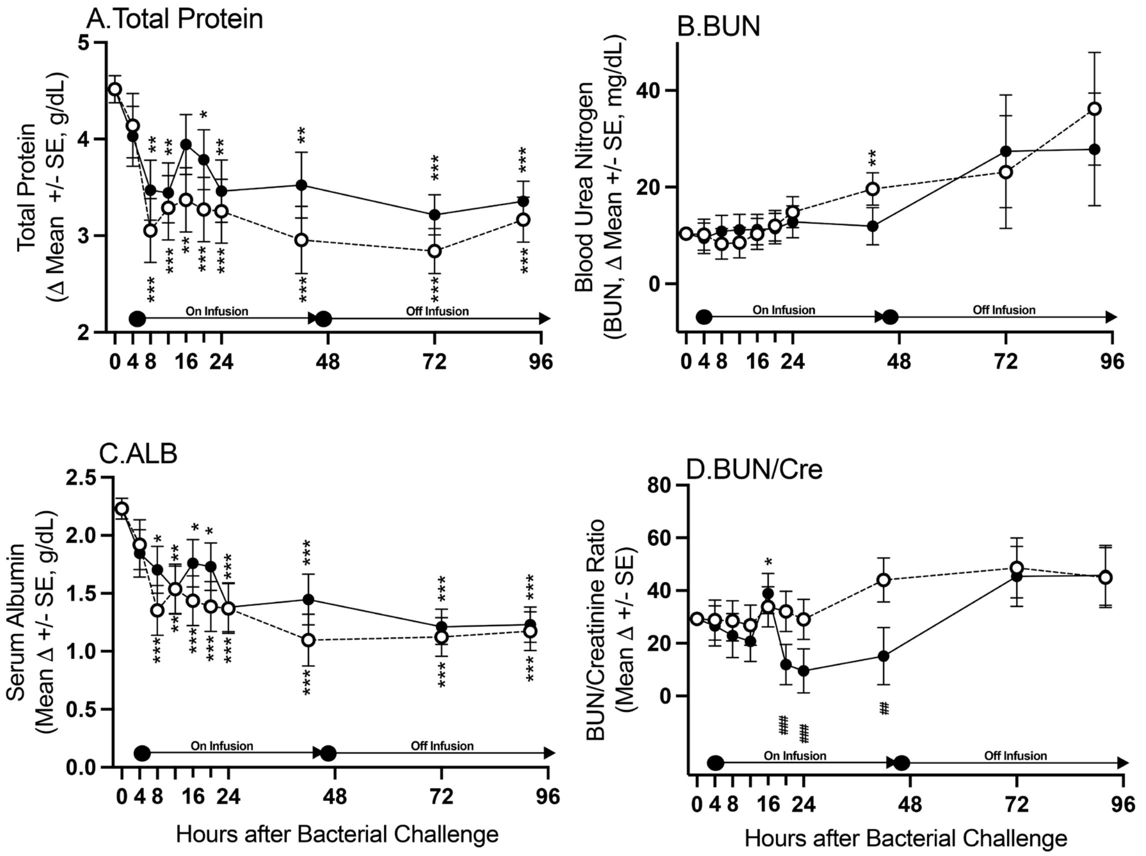


P value
Change from Baseline (T0)
* p : 0.01 - 0.05
** p : 0.001 - 0.01
*** p < 0.001

P-Value
Effect of Epinephrine
#p: 0.01-0.05
##p: 0.001-0.01
###p < 0.001

**e-supplementary figure 3: The format is similar to figure 1**

**e-supplementary figure 4**

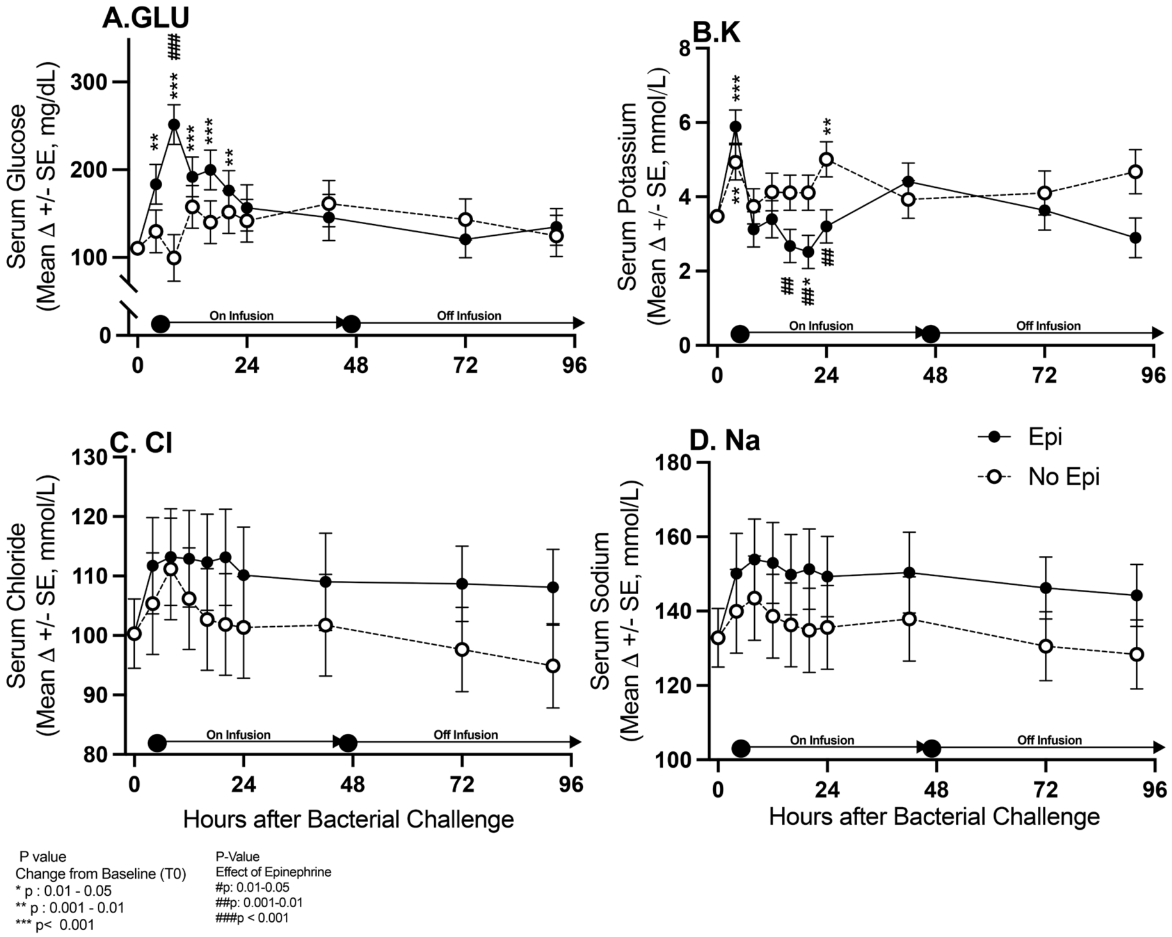


**e-supplementary figure 4: The format is similar to figure 1**

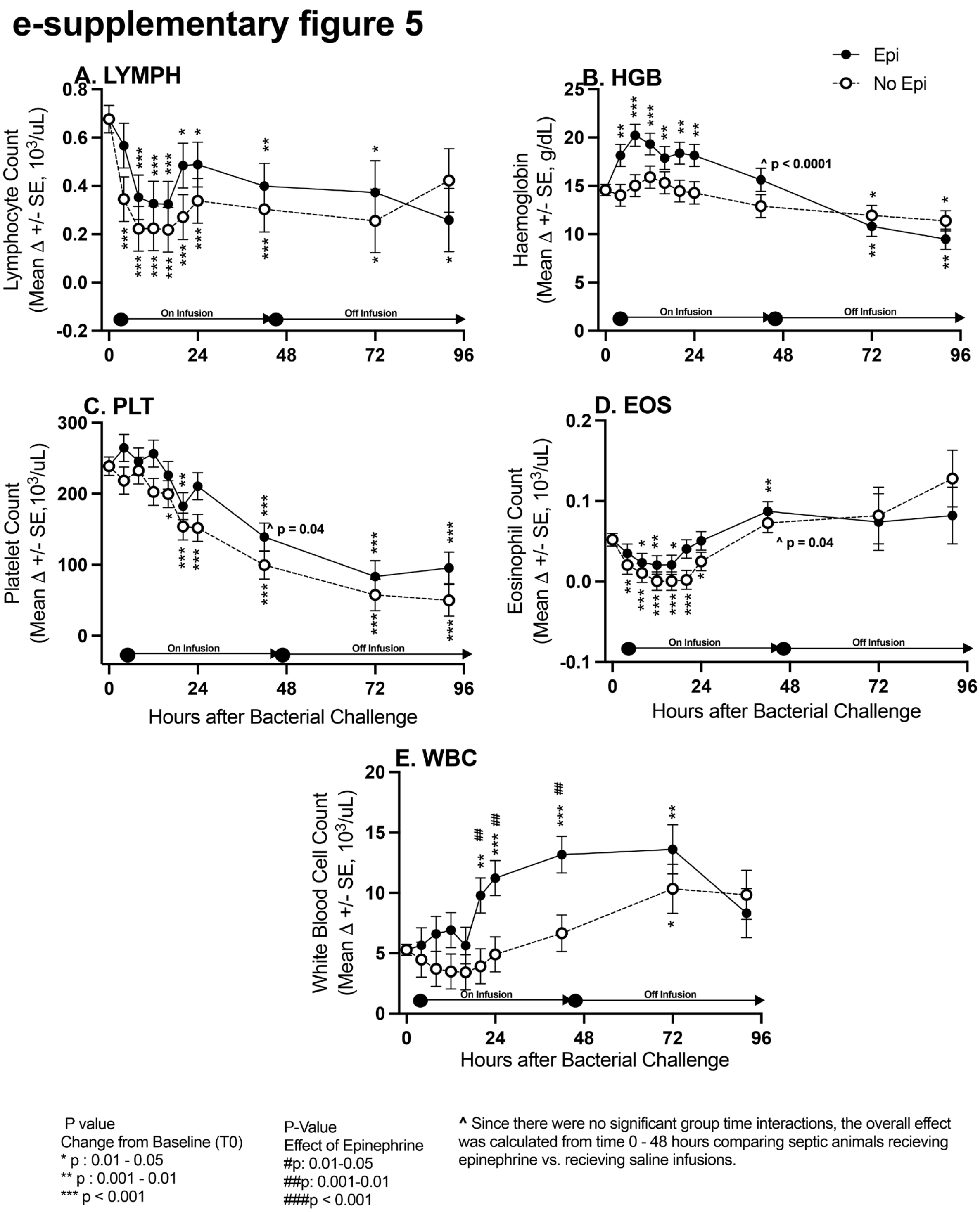


**e-supplementary figure 5: The format is similar to figure 1**
